# Microfluidic platform for microbial spore germination studies in multiple growth conditions

**DOI:** 10.1101/2024.05.13.593863

**Authors:** Léa S. Bernier, Aislinn Estoppey, Saskia Bindschedler, Guy-Bart Stan, Pilar Junier, Claire E. Stanley

## Abstract

Spores are highly resistant dormant cells, adapted for survival and dispersal, that can withstand unfavourable environmental conditions for extended periods of time and later reactivate. Understanding the germination process of microbial spores is important in numerous areas including agriculture, food safety and health, and other sectors of biotechnology. Microfluidics combined with high-resolution microscopy allows to study spore germination at the single-cell level, revealing behaviours that would be hidden in standard population-level studies. Here, we present a microfluidic platform for germination studies where spores are confined to monolayers inside microchambers, allowing the testing of four growth conditions in parallel. This platform can be used with multiple species, including non-model organisms, and is compatible with existing image analysis software. In this study, we focused on three soil dwellers, two prokaryotes and one fungus, and revealed new insights into their germination. We studied endospores of the model bacterium *Bacillus subtilis* and demonstrated a correlation between spore density and germination in rich media. We then investigated the germination of the obligate-oxalotrophic environmental bacterium *Ammoniphilus oxalaticus* in a concentration gradient of potassium oxalate, showing that lower concentrations result in more spores germinating compared to higher concentrations. We also used this microfluidic platform to study the soil beneficial filamentous fungus *Trichoderma rossicum*, showing for the first time that the size of the spores and hyphae increase in response to increased nutrient availability, while germination times remain the same. Our platform allows to better understand microbial behaviour at the single-cell level, under a variety of controlled conditions.

**One-Sentence Summary:** A microfluidic platform developed for spore germination studies in multiple growth conditions provides new insights into the germination of spores at the cellular level from three soil dwellers.

## Introduction

Dormant cells have reduced metabolic activity and increased resistance to unfavourable environments (McDonald *et al*. 2023), constituting an important survival strategy for many microorganisms (Jones and Lennon 2010). Dormant cells survive harsh environmental conditions and the absence of nutrients, having the potential to reactivate once conditions improve. Hence, dormancy constitutes a strategy for species to survive despite the variability of their environment over time. These dormant cells include spores, but also hypnozygotes, cysts, oospores, akinetes and ephippia, for example (Mestre and Höfer 2021). The best studied bacterial spores are endospores, formed by *Bacillales* and *Clostridiales* species (Setlow *et al*., 2017). Unlike bacteria, fungi can produce a large variety of spores, even within a single species, and their spores vary greatly in size and function. For example, *Trichoderma* species make both asexual conidia (3-5 µm) and resting chlamydospores (8-10 µm) (Lewis and Papavizas 1983). In contrast, spores of arbuscular mycorrhizal fungi (AMF) species typically measure 50-750 µm, and sometimes over 1 mm (Walker 2013). In addition to the resistance to extreme environmental conditions, dormancy facilitates dispersal, as resistant dormant cells remain viable for longer periods of time in adverse conditions, and hence survive travel or transport. Fungal spores, for example, have been reported to travel thousands of kilometres over tropical and subtropical oceans (Mayol *et al*. 2017).

Understanding how, when, and why spores germinate is relevant to many fields, from agriculture to health and food. Spores have the potential to contribute to sustainable agriculture via their extensive use in the formulation of biocontrol agents (Martinez *et al*. 2023; Poulaki and Tjamos 2023; Ye *et al*. 2023). Indeed, their resistant dormant nature provides an easy way to facilitate storage and prolong shelf life. However, germination should be precisely understood so it can be prevented until the biocontrol agent is in soil and ensured afterwards. Many pathogens produce spores as a mechanism of survival in the absence of a suitable host or as a dispersal mechanism during disease expansion. Spores also form a critical component of microbiomes and can contribute to resistance and relapse in human pathology (Tetz and Tetz 2017). They have also found use as biotechnological tools, with applications ranging from vaccine delivery to components of functional materials such as self-healing concrete (Paul *et al*. 2019; Zhang *et al*. 2020).

Spore germination is the transition phase in the life cycle of spore-formers, where a spore reactivates vegetative growth. Fungal spore germination typically involves active swelling (rehydration) and cell wall remodelling, followed by the formation of a germ tube and filamentous growth (Osherov and May 2006). In endospore-forming bacteria, such as the well-known model bacterium *Bacillus subtilis*, this process can be characterised by monitoring the transition from a phase-bright spore to a phase-dark vegetative cell. This change is accompanied by rehydration, release of dipicolinic acid (DPA), cortex hydrolysis and coat disassembly (Zhou *et al*. 2022). Bacterial spore germination studies rely on many different techniques to measure this process, often used in conjunction with one another. Some of the oldest techniques that are still used to this day involve measuring the loss of turbidity of media (Fleming and Ordal 1964; Wax, Freese and Cashel 1967; Prasad 1974; Yoko Yasuda 1984; Alzahrani and Moir 2014; Krajčíková, Bugárová and Barák 2021) or counting of colony forming units (CFU) (Setlow, Cowan and Setlow 2003; Nagler *et al*. 2014). These measurements are taken at the population level and do not provide information about potential cell-to-cell variability. Additionally, these techniques are indirect, and measure not only germination itself, but also the vegetative cell growth that follows. However, they are very well known and do not require any specialist equipment. Another common population-level method applicable in the case of endospore-forming bacteria is the measurement of DPA release (Setlow, Cowan and Setlow 2003; Black *et al*. 2005; Yi and Setlow 2010; Butzin *et al*. 2012; Grela *et al*. 2018). DPA release can be monitored in real-time using fluorescence spectroscopy in standard plate reader assays thanks to the addition of terbium chloride, which results in the formation of a Tb^3+^-DPA complex (Yi and Setlow 2010). This automated process allows to easily monitor DPA release for large numbers of samples simultaneously but remains a population-level method. Other methods are necessary to reveal variability at the single spore level. They include flow cytometry (Black *et al*. 2005; Saggese *et al*. 2022), Raman spectroscopy (Wang *et al*. 2011; He *et al*. 2018; Wu *et al*. 2021), and various microscopy techniques such as differential interference contrast (DIC) microscopy (Wang *et al*. 2011; Butzin *et al*. 2012; Nagler *et al*. 2014; He *et al*. 2018), transmission electron microscopy (TEM) (Lu *et al*. 2021; Tsugukuni *et al*. 2021), fluorescence microscopy (Thrane *et al*. 2000; Mutlu *et al*. 2018; Tsugukuni *et al*. 2021), and phase-contrast microscopy (Vepachedu and Setlow 2004; Løvdal *et al*. 2012; Nagler *et al*. 2014; Wu *et al*. 2021; Kikuchi *et al*. 2022). Flow cytometry allows for the monitoring of a very high number of single cells but cannot capture the progression of germination in the same cell over time. Raman spectroscopy has the advantage of providing label-free molecular-level single-cell insights but requires expensive specialist equipment and skills. Microscopy techniques on the other hand allow for single-cell level long-term monitoring of germination events and outgrowth, and, although they can be expensive, microscopes are commonly found in microbiology laboratories (Trunet, Carlin and Coroller 2017).

In recent years, microfluidic or so-called Spore-on-a-Chip technologies have emerged as a powerful tool for the study of spores (Bernier *et al*. 2022). Microfluidic platforms are tools that allow handling of fluids at the microscale. As such, miniaturisation leads to lower volume requirements, enhanced throughput, and parallelisation, ultimately leading to cost savings and improved performance (Cottet and Renaud 2021). More importantly, microfluidic technologies provide a powerful approach for single-cell level studies. Confinement of individual cells to a single monolayer allows each cell to be tracked with high spatial and temporal resolution, thus revealing behaviours that remain hidden at the population level, such as the presence of dormant sub-populations and population heterogeneity (Arvaniti and Skandamis 2022; Shan *et al*. 2023).

Only a handful of microfluidic platforms have been applied to bacterial and fungal spore germination studies previously. Some rely on electrical measurements to allow real-time monitoring of germination at a small population level. For example, Zabrocka *et al*. developed a microfluidic device that measures time-dependent changes in the electrical conductivity of a nanofibre network to detect *B. subtilis* germination (Zabrocka *et al*. 2015), and Liu *et al*. used electrodes for automatic electrical detection of *Bacillus anthracis* germination (Liu *et al*. 2007). To achieve single-cell level data, some studies have explored small culture chambers with media perfusion. These culture chambers have been used to monitor fungal spore germination using a single chamber (Grünberger *et al*. 2017), or a small array of culture chambers together on one device (Barkal *et al*. 2016) (*Penicillium chrysogenum* and *Cryptococcus neoformans,* respectively). The microfluidic single-cell cultivation (MSCC) device employs arrays of microchambers, allowing analysis of the germination of the filamentous bacterium *Streptomyces lividans* under multiple media compositions, which revealed surprising stability in behaviour despite previous observations in large-scale cultures (Koepff *et al*. 2018). For AMF spores, Richter et al. have recently presented the *AMF-SporeChip*, which revealed new insights into AMF spore germination and hyphal growth (Richter *et al*. 2024).

We developed a microfluidic platform for the study of microbial spore germination in four different growth conditions in parallel, all included within a single device. This platform is compatible with the study of multiple growth conditions, such as pH, water potential (polyethylene glycol (PEG)), salinity, metal stress and trophic factors. Here, we focused on the latter and varied the media composition. It was designed to be easy to use and adaptable to different microorganisms and research questions. Cells are constrained to a single monolayer to facilitate image analysis. In this study, we focused on three spore-forming soil dwellers already used, or with the potential to be used, as biocontrol agents. First, we quantified germination of the model organism *B. subtilis* upon exposure to (i) four different growth conditions and (ii) a concentration gradient of the well-known germinant L-alanine and investigated the effect of spore density on germination. L-alanine has been described as the “most universal germinant for spores from a diversity of origins” (Moriyama *et al*. 1996). Its role in the initiation of *B. subtilis* spore germination has been studied for over 50 years (Wax, Freese and Cashel 1967). We then studied the environmental endospore-forming soil bacterium *Ammoniphilus oxalaticus* (a non-model organism), evaluating the effect of potassium oxalate on germination with high temporal and spatial resolution and in a quantitative manner. Finally, we investigated the effect of nutrient availability on the germination of asexual spores (conidia) from the soil beneficial filamentous fungus *Trichoderma rossicum*. These experiments showed insights into (i) the effect of spore density on the germination of *B. subtilis*, showing that higher spore density leads to faster germination in rich media, (ii) the influence of potassium oxalate on *A. oxalaticus* spore germination, where a higher percentage of spores were found to germinate at lower concentrations of potassium oxalate than at higher concentrations, as well as (iii) the response of *T. rossicum* to varying levels of nutrients, demonstrating that increased nutrients availability leads to larger hyphae, but does not affect the time of germination.

## Materials and methods

### Pre-cultivation, media and bacterial strains

*Bacillus subtilis* (CotY8GA) CotY-GFP (Imamura *et al*. 2011) spores were prepared as follows. Nutrient agar plate cultures consisting of 28 g L^-1^ nutrient agar powder (Oxoid) supplemented with 25 µg mL^-1^ chloramphenicol (Sigma) were incubated at 30 °C for two weeks. Spores were harvested and purified as previously described (Amon *et al*. 2020). Briefly, the spores were scraped off the plates with autoclaved milliQ water, heat treated in an oven at 70 °C for 30 minutes and washed three times (4000 rpm centrifugation for 5 minutes) with autoclaved milliQ water. The pellet was then resuspended in 20% (w/v) HistoDenz (Sigma) and incubated on ice for 30 minutes. This suspension was then gently pipetted onto a 2-step HistoDenz gradient (40% Histodenz layered on 50% Histodenz) and centrifuged at 4000 rpm for 90 minutes. After discarding the supernatant, the pellet was washed (4000 rpm centrifugation for 5 minutes) 3 times with autoclaved milliQ water, resuspended in autoclaved milliQ water and the spore suspensions were stored at 4 °C. The spore suspensions were checked by microscopy and diluted to an OD 600 of 0.1 before use (approximately 1 × 10^7^ spores per mL).

The synthetic media (SM) used as the base for the different growth conditions for the *B. subtilis* experiments contained 1 mM fructose, 1 mM glucose and 10 mM potassium chloride. The four conditions in the first experiment were 13 g L^-1^ nutrient broth (NB) (Oxoid), SM with 10 mM L-alanine (Sigma), SM with 10 mM D-alanine (Sigma), and SM. To obtain the L-alanine concentration gradient solutions, SM was supplemented with L-alanine at 1 µM, 100 µM, 10 mM and 1 M concentrations. All solutions were sterile-filtered using 0.2 µm filters (Minisart NML, Sartorius).

*Ammoniphilus oxalaticus* HJ (Estoppey 2023) spores were prepared as follows. *A. oxalaticus* cells were cultured on bilayered Schlegel AB media plates supplemented with 4 g L^-1^ calcium oxalate (Aragno and Schlegel 1992). The first layer consisted of Schlegel AB agar and the second layer of Schlegel AB agar supplemented with calcium oxalate, in a 2:1 ratio. This media was prepared following the method from medium 81 from (Koblitz *et al*. 2023). Briefly, Schlegel A contained 9 g L^-1^ Na_2_HPO_4_.12H_2_O (M-152A), 1.5 g L^-1^ KH_2_PO_4_ (M-81), 1 g L^-1^ NH_4_Cl (M-194), 0.2 g L^-1^ MgSO_4_.7H_2_O (M-122A), 1 mL L^-1^ SL-6 10x Trace elements solution (see medium 27 from (Koblitz *et al*. 2023)), while Schlegel B contained 0.92 g L^-1^ Fe(NH_4_)(SO_4_)_2_.12H_2_O (M-61) and 1.12 g L^-1^ CaCl_2_.2H_2_O (M-25). Schlegel AB contains Schlegel A and 1 % Schlegel B, and Schlegel AB agar has a technical agar concentration of 15 g L^-1^. The plates were incubated at room temperature for eight weeks. Spores were harvested from each plate using 1 mL of physiological water (autoclaved milliQ water with 0.9% NaCl) and used immediately. Liquid Schlegel AB media supplemented with 0.3 g L^-1^, 0.65 g L^-1^, 0.4 g L^-1^ and 8 g L^-1^ potassium oxalate was used for germination experiments. In the control experiment, the four conditions were Schlegel AB with 0.65 g L^-1^ potassium oxalate, Schlegel AB only (0 g L^-1^ potassium oxalate), NB and physiological water. All solutions were sterile-filtered using 0.2 µm filters (Minisart NML, Sartorius). We measured the pH of Schlegel AB with and without potassium oxalate using pH 5.0 – 10.0 pH indicator strips (MQuant, Supelco).

*Trichoderma rossicum* NEU135 (Bravo *et al*. 2013) spores were prepared as follows. Plate cultures composed of 39 g L^-1^ potato dextrose agar (PDA) (Difco Potato dextrose agar, Becton Dickinson) were incubated at 26 °C for 5 weeks. Spores were harvested from the surface of each plate with 2 mL of autoclaved milliQ water and filtered through autoclaved glass wool. After centrifugation for 1 minute at 2000 rpm, the supernatant containing purified spores (as checked by microscopy) was used for experiments. Potato dextrose broth (PDB) (Difco Potato dextrose broth, Becton Dickinson) diluted with autoclaved milliQ water was used for the 100% (24 g L^-1^), 1% and 0.1% PDB conditions. The control (0% PDB solution) consisted of physiological water (autoclaved milliQ water with 0.9% NaCl). All solutions were also sterile-filtered using 0.2 µm filters (Minisart NML, Sartorius) for experiments.

### Microfluidic device fabrication

The design of the device was constructed in AutoCAD (Autodesk) and printed onto a photolithography mask (Micro Lithography Services Ltd., UK). It included 324 cultivation chambers arranged in 12 arrays consisting of 27 open box chambers (90 µm × 80 µm) (Täuber *et al*. 2020) to which we added 4 channels (each 100 µm in width) leading to 4 inlets. Thanks to laminar flow, these 4 inlets resulted in 4 zones in the device, each spanning 3 arrays. There were hence 81 chambers in each condition. The first layer was etched in the 4-inch silicon wafer (Diameter: 4”; Orientation: <100>; Dopant: P(Boron); Resistivity: 0-100 ohm cm; Centre thickness: 425-550 μm; Surface: single side polished, Si-Mat Silicon Materials). First, a layer of SU-8 2002 (Microchem) was spin coated at 6000 rpm (ramp 500 rpm/s) for 30 seconds (Spin Coater WS-650MZ Modular, Laurell Technologies Corporation) onto a 100 mm diameter silicon wafer. After the soft bake (95 °C for 900 seconds), the wafer was exposed in a UV-KUB 3 mask aligner with an energy dose of 90 mJ cm^-^² (Kloe). After the post-exposure bake (95 °C for 900 seconds), it was developed in SU-8 developer (1,2-propanediol monomethyl ether acetate, ThermoScientific) rinsed with isopropyl alcohol and air dried. Then, the pattern was etched into the silicon wafer using the PlasmaPro 100 Cobra ICP Etching chamber of the Oxford Instruments Plasma Cluster to achieve a height of 0.73 ± 0.01 µm. The wafer was etched for 5 minutes, followed by oxygen plasma cleaning (Oxford Instruments) for 5 minutes. The wafer was sonicated in milliQ water for 5 minutes. To form the second layer, a 10 µm high layer of SU-8 2010 photoresist (Kayaku Advanced Materials) was spin coated at 3500 rpm (ramp 500 rpm/s) for 45 seconds (Spin Coater WS-650MZ Modular, Laurell Technologies Corporation). After the soft bake (95 °C for 150 seconds), the wafer was exposed to ultra-violet light in the UV-KUB 3 mask aligner with an energy dose of 140 mJ/cm². After the post-exposure bake (95 °C for 210 seconds), it was developed in SU-8 developer, rinsed with isopropyl alcohol and air dried. Finally, it was hard baked for 15 minutes at 150 °C. The wafer was silanised under vacuum for 2 hours with 50 μL chlorotrimethylsilane (Sigma).

Poly(dimethylsiloxane) (PDMS) (Sylgard 184, Dow Corning) was prepared at a 10:1 ratio of base to curing agent and degassed for 1 hour under vacuum in a desiccator. It was poured onto the master mould and cured overnight at 70 °C. It was then removed from the mould and cut into individual pieces. Subsequently, the 4 inlets and the outlet were punched with a precision cutter (Ø 1.02 mm; Syneo, USA). After washing and drying, the PDMS slabs were bonded onto glass-bottomed petri dishes (Ø dish = 35 mm, Ø glass = 23 mm, glass thickness = 0.17 mm; Fluorodish, World Precision Instruments) using a Zepto plasma cleaner (Diener Electronic; vacuum pressure 0.75 mbar, power 50%, coating time 1 minute).

### Setup and microfluidic cultivation

The spores were loaded into the device as described previously (Täuber *et al*. 2020). Briefly, approximately 100 µL of the spore suspension was introduced into the device from the outlet by hand. Random flow patterns were generated, and together with the flexibility of PDMS, allowed spores to enter the microchambers. They remain trapped inside the shallow chamber and form a monolayer. This loading method results in a random distribution of spores across the chambers. The syringe (1 mL Henke-Ject Luer, Henke Sass Wolf) was then disconnected from the outlet, and the device inlets connected to 4 syringes (5 mL or 10 mL Luer Lock Henke-Ject, Henke Sass Wolf) containing various types of growth media, as well as a piece of Tygon tubing (inner diameter 0.030 inches, outer diameter 0.090 inches, Cole-Parmer, US) connected to a waste container. Each piece of Tygon tubing was connected to the device via a metal pin (stainless steel catheter coupler, 20 ga x 15 mm, Linton Instrumentation) and to the syringe via a blunt-end luer lock syringe needle (Darwin Microfluidics). The tubings were always connected starting from the lower concentration (or negative control) to limit cross-contamination within the different zones of the device. All tubings, pins, and blunt-end luer lock syringe needles were autoclaved before use. A 6-channel programmable syringe pump (AL-1600, World Precision Instruments) was used to achieve a flow rate of 5 µL per minute throughout the experiments.

To characterise the microfluidic device, a concentration gradient of milliQ water with fluorescein at 0, 0.025, 0.125 and 0.25% (w/vol) was generated. All solutions were filtered using 0.2 µm filters (Minisart NML, Sartorius).

### Live cell imaging

Timelapse images of the device were acquired with an inverted microscope (Eclipse Ti-2, Nikon), equipped with an oil immersion ×100/1.3 NA (numerical aperture) Plan Fluor objective lens and connected to a camera (DS-Qi2, Nikon). The stage incubator was set to 30 °C for the experiments with *B. subtilis* and 26 °C for the experiments with *A. oxalaticus* and *T. rossicum*. A single field of view corresponds to 1 chamber in the device, and 4 positions (4 chambers) were manually selected in each zone (i.e., for each condition) for bacterial spore germination experiments, and 5 positions for fungal spore germination experiments. We selected chambers with spores that were – when possible – well separated (i.e., not in clusters) so that the images would be easier to analyse. The timelapses were taken for the bacterial and fungal spore germination experiments with an interval of 5 and 60 minutes, respectively. The experiments were run for 5 hours for *B. subtilis,* and 20 hours for *A. oxalaticus* and *T. rossicum*. The perfect focus system (PFS) was used with the NIS-Elements Advanced Research imaging software (Nikon).

### Image and Data Analyses

Images were processed and analysed using Fiji (Schindelin *et al*. 2012). The bacterial spore germination time was assessed with the aid of the ImageJ plugin SporeTrackerX (Omardien *et al*. 2018). The brightness threshold was set manually for each chamber, and each timelapse was checked manually to differentiate non-germinating spores from image artefacts. For fungal spore germination experiments, the time of apparition of the first germ tube was taken as the time of the frame preceding its apparition. The maximum swelling diameter of the spore, as well as the diameter of the lead hypha, were measured using Fiji.

For *B. subtilis*, 1-20 spores were analysed in each chamber for the experiment with four growth conditions, and 0-14 spores per chamber were analysed for the L-alanine gradient experiment. Spores were prepared in advance at high concentrations which allowed higher numbers of spores in the *B. subtilis* experiments. On the other hand, spores were harvested immediately prior to experiments from a single plate for experiments with *A. oxalaticus* and *T. rossicum,* resulting in lower spore counts. In the *A. oxalaticus* experiment, 1-5 spores were analysed in each chamber, and 1-4 spores for *T. rossicum*.

All graphs and statistics were produced in GraphPad Prism version 10.0.3 (GraphPad Software). The frequency distribution was calculated as the relative frequency (fractions) of germinating spores within each 5-minute interval, i.e., what fraction of analysed spores germinated during each 5-minute interval. It was calculated separately for each condition. All boxplots were produced with boxes extending from the 25^th^ to 75^th^ percentiles, whiskers going from minimum to maximum values, and a line displaying the median. Significance was tested using Mann Whitney tests to compare each pair of conditions. Spearman correlations were calculated between the number of spores per chamber and the germination times. We used a threshold significance level of 0.05 and considered results to be extremely significant (****) when p < 0.0001 and (***) when 0.0001 < p > 0.001, very significant (**) when 0.001 < p > 0.01, significant (*) when 0.01 < p > 0.05, and not significant (ns) when p ≥ 0.05. The bar charts were produced by taking the percentage of all analysed spores that germinated or remained dormant (non-germinated) at the end of the experiments.

## Results

### Microfluidic device design and operation

We developed the 4-conditions microfluidic chemostat (4CMC, Figure 1A) to culture microbial spores in microchambers under four different growth conditions simultaneously within one single device. This device design makes it possible to study germination at the cellular level for different organisms. The 4CMC houses 324 cultivation chambers (Figure 1B) (Täuber *et al*. 2020) arranged in 12 arrays consisting of 27 open box chambers (90 µm x 80 µm) that have a height of 0.73 ± 0.01 µm to ensure spore entrapment (Figure 1C) and subsequent monolayer growth, while the rest of the device has a height of 10 µm. Spore entrapment is made possible by the combination of random flow patterns introduced during loading and the flexibility of PDMS. The spores are manually loaded into the device, resulting in a random distribution of spores inside chambers and therefore generating a range of spore occupancies. The 4CMC design includes four medium inlet channels to allow up to four different growth conditions to be studied in parallel in the same device. As a result of operating under a laminar flow regime, different zones within the device can be established and correspond to the four growth conditions to be evaluated. The transfer of nutrients inside the cultivation chambers occurs via diffusion only (Täuber *et al*. 2020). Each zone spans 3 arrays, and the different zones were characterised using fluorescein experiments (Figure 1D) as well as COMSOL simulations (Supplementary Figure S1). Figure 1E shows an example of endospore germination as observed within a chamber in the 4CMC device.

**Figure 1:**
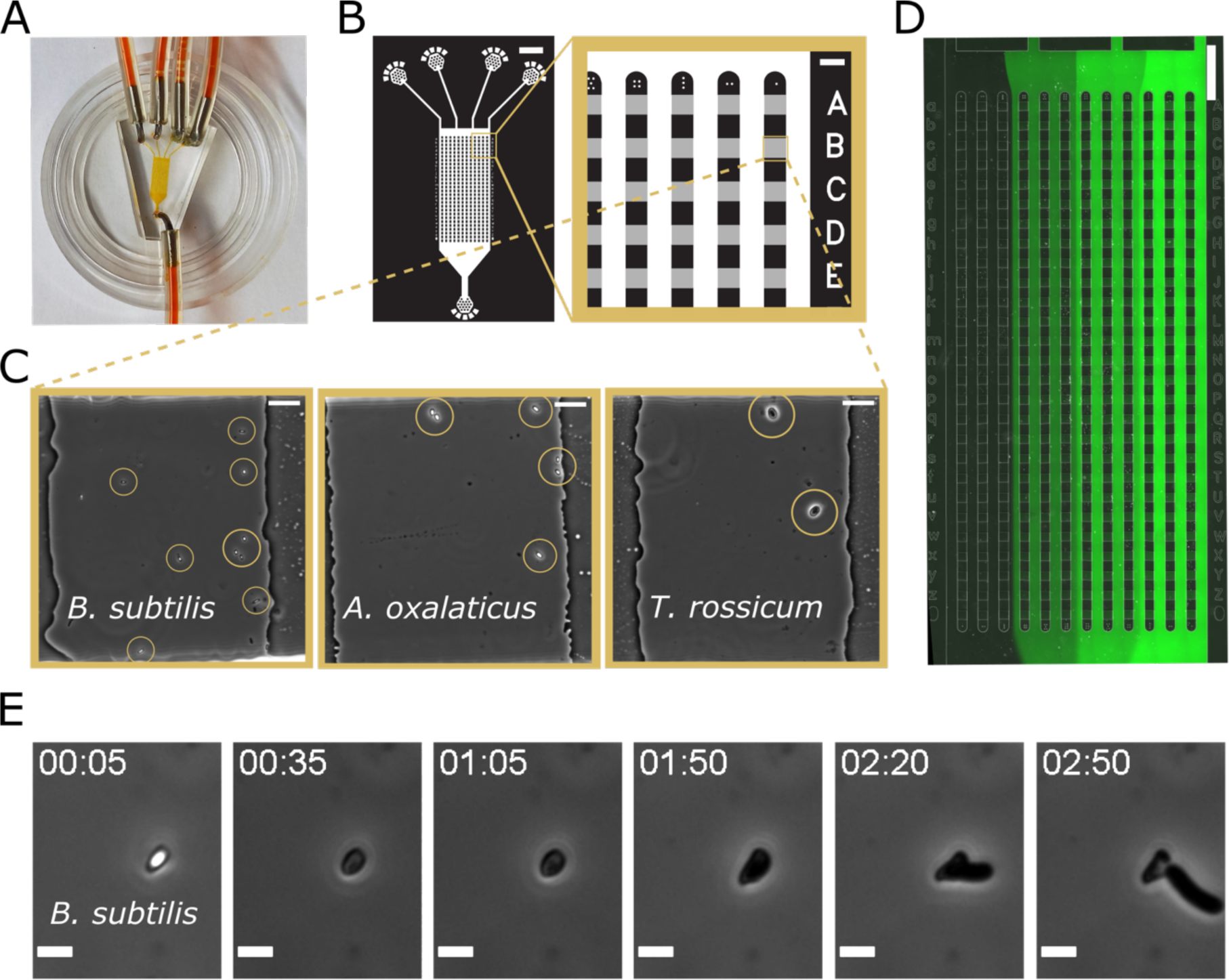
Design of the 4-conditions microfluidic chemostat (4CMC) device for cultivation of microcolonies under four growth conditions. **(A)** Photograph of the microfluidic device inside a 35 mm glass-bottomed Petri dish. **(B)** Design of the microfluidic device including the arrays of cultivation chambers. The cultivation chambers (in grey) have a height of approximately 750 nm to ensure monolayer growth while the supply channels (in white) are 10 µm high. Scale bars, 1 mm (left) and 100 µm (right). **(C)** Examples of individual chambers containing, from left to right: *Bacillus subtilis, Ammoniphilus oxalaticus,* and *Trichoderma rossicum* spores. Scale bars, 10 µm. **(D)** Flow profile within the microfluidic device used with a concentration gradient of fluorescein solutions. Scale bar, 500 µm. **(E)** Phase contrast images of *B. subtilis* cell in the microfluidic device at different time points showing the phase bright-spore (00:05) turning phase-dark (from 00:35), then outgrowth (from 01:50) and cell detachment from the spore’s coat (02:50). Scale bar, 2 μm. Time stamp format, hh:mm.

### Responsiveness of *B. subtilis* spores to different germinants

In the model organism *B. subtilis*, the respective initiation and inhibition effect of L- and D-alanine on germination, and their mechanism of action are well defined (Yoko Yasuda 1984; Yasuda and Tochikubo 1985; McCann *et al*. 1996; Løvdal *et al*. 2012). Hence, in the first set of experiments, the 4CMC device was used to study the germination of *B. subtilis* spores upon exposure to the following media conditions: rich media (nutrient broth, NB), synthetic media (SM) supplemented with L-alanine (10 mM), SM supplemented with D-alanine (10 mM), and SM without supplementation (Video 1). Germination events were analysed using SporeTrackerX (Omardien *et al*. 2018). The software defines germination events according to a drop in brightness, which is assumed to correspond to the uptake of water and release of DPA (Pandey *et al*. 2013). The relative frequency distributions of the spore germination time, which show what fraction of spores germinated within each 5-minute interval (as obtained from the time-lapse experiments), are displayed in Figure 2A-D. Figure 2E shows a comparison of the germination times in each condition. The overall percentage of spores that germinated or remained as spores (non-germinated) after 5 hours is displayed in Figure 2F. In NB and SM + L-alanine all spores analysed germinated, but germination was faster in NB as compared to SM + L-alanine. In NB, all spores germinated within 1 hour, and over 90% germinated within 15 minutes, whereas in SM + L-alanine, a smaller fraction (under 70%) of the analysed spores germinated within the first 15 minutes and the rest took up to 2 hours to germinate. On the contrary, very few of the tracked spores germinated in SM supplemented with D-alanine (7.4%) and SM alone (4.4%). The few spores that did germinate in SM + D-alanine did so over the course of the entire experiment, some germinating within the first 15 minutes, but others after 1, 2 or even 3 hours. In contrast, the ones that germinated in SM alone (4.4%) did so within the first 15 minutes. Additionally, we found a negative correlation (Spearman’s rank correlation coefficient r = -0.8) between the number of spores per chamber and their germination time in NB (but no correlation in SM + L-alanine) (Figure 2G-H); indeed, the more spores in a chamber, the faster they germinated. The number of spores per chamber showed a positive correlation with the number of germinating spores in NB and SM + L-alanine, and a positive correlation with the number of non-germinating spores in SM + D-alanine and SM alone (Supplementary Figure S2). Mann Whitney tests showed statistically significant differences between the germination times in all conditions with the exception of SM (Figure 2E).

**Figure 2:**
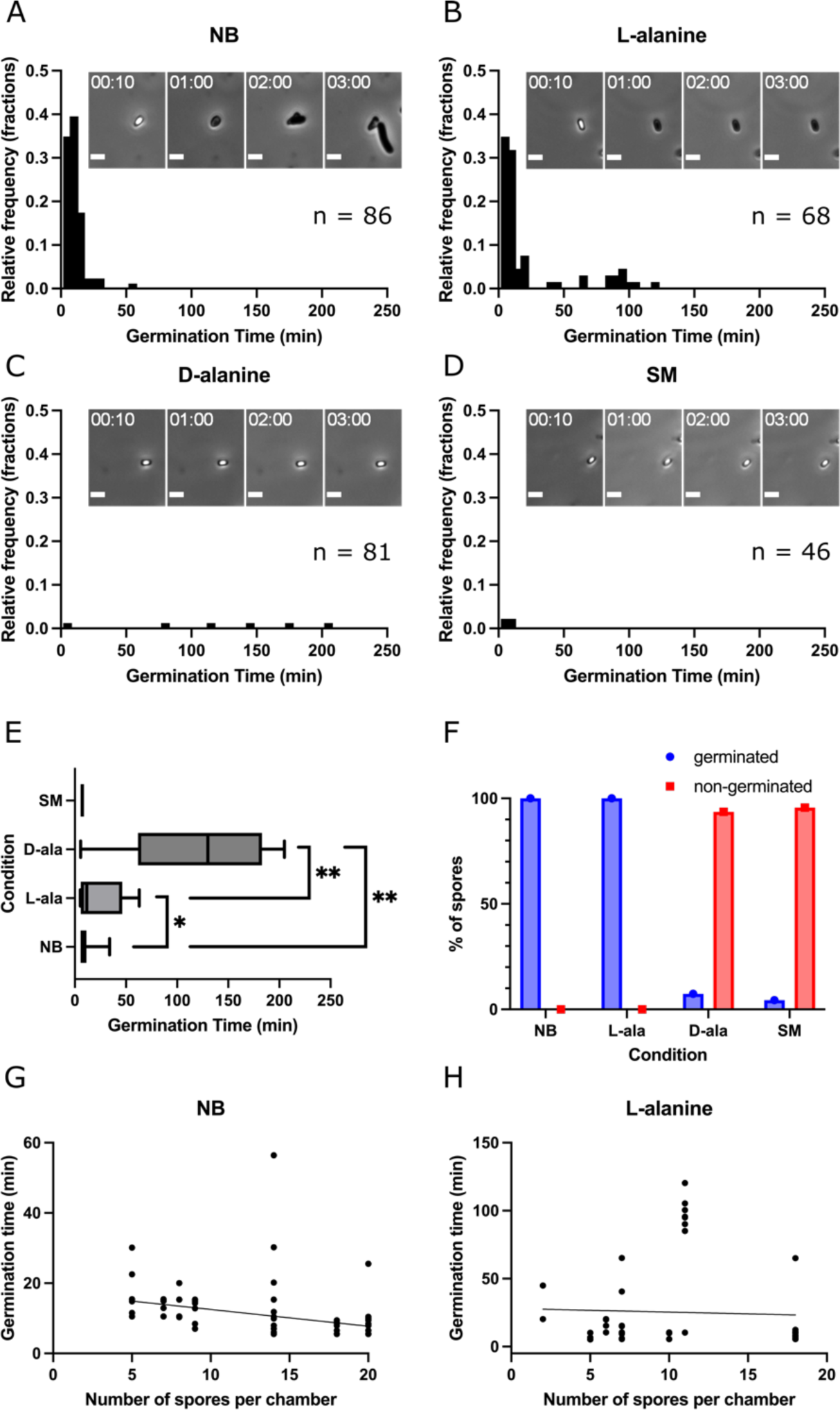
Germination of *Bacillus subtilis* in four different media conditions applied simultaneously in the 4CMC device. Example timelapses for each condition available in Video 1. **(A-D)** Relative frequency distributions of the time of germination in each condition. The graphs show the proportion (i.e., relative frequency) of the analysed spores germinated within 5-minute intervals. The total number of analysed spores (n) in each condition is indicated. Phase contrast microscope images of *B. subtilis* in the microfluidic device over time showing an individual spore. Scale bars, 2 µm. Time stamp format, hh:mm. **(E)** Boxplot of the germination times in each condition. The line shows the median, the box extends from the 25^th^ to 75^th^ percentiles and the whiskers from min to max. Mann Whitney tests showed statistically significant differences (*p* < 0.05) between each condition except for SM. **(F)** Bar chart showing the percentage of spores that germinated or remained dormant (non-germinated) after 5 hours of incubation for the four conditions compared within the microfluidic device. **(G)** Germination time of individual spores in relation to number of spores per chamber in rich media (NB), showing a negative correlation (Spearman r = -0.8) between the germination times and the number of spores per chamber. Data from 8 chambers were analysed; note, two chambers contained 5 spores per chamber. **(H)** Germination times in relation to number of spores per chamber in L-alanine, showing no correlation (Spearman r = 0.06) between the germination times and the number of spores per chamber. Data from 8 chambers were analysed; note, two chambers contained 7 spores per chamber.

The 4CMC device was then used to study the germination of *B. subtilis* spores in more detail upon exposure to a concentration gradient (1 μM, 100 μM, 10 mM and 1 M) of L-alanine (Video 2). L-alanine has been studied extensively, but not using methods that give direct single spore level information about germination events (Lyu *et al*. 2023). The relative frequency distributions of the spore germination time, showing what fraction of the analysed spores germinated within each 5-minute interval, are displayed in Figure 3A-D. Figure 3E compares the germination times in each condition. All the spores that germinated did so within 25 minutes in all conditions, except for a single spore that germinated after 4 hours and 50 minutes (in the 100 µM condition). Mann Whitney tests showed no statistically significant differences between the germination times in each condition (Figure 3E). The 10 mM L-alanine conditions in both sets of experiments have median germination times of 11.6 and 10.5 minutes, respectively, and no statistically significant difference as shown in Supplementary Figure S3. The overall percentage of spores that germinated or remained as spores (non-germinated) after 5 hours is displayed in Figure 3F. Only 33.3% of analysed spores germinated at 1 µM concentration, but already 90.9% germinated at 100 µM concentration. This percentage reached 97.1% at 10 mM concentration, and all the analysed spores germinated at 1 M concentration.

**Figure 3:**
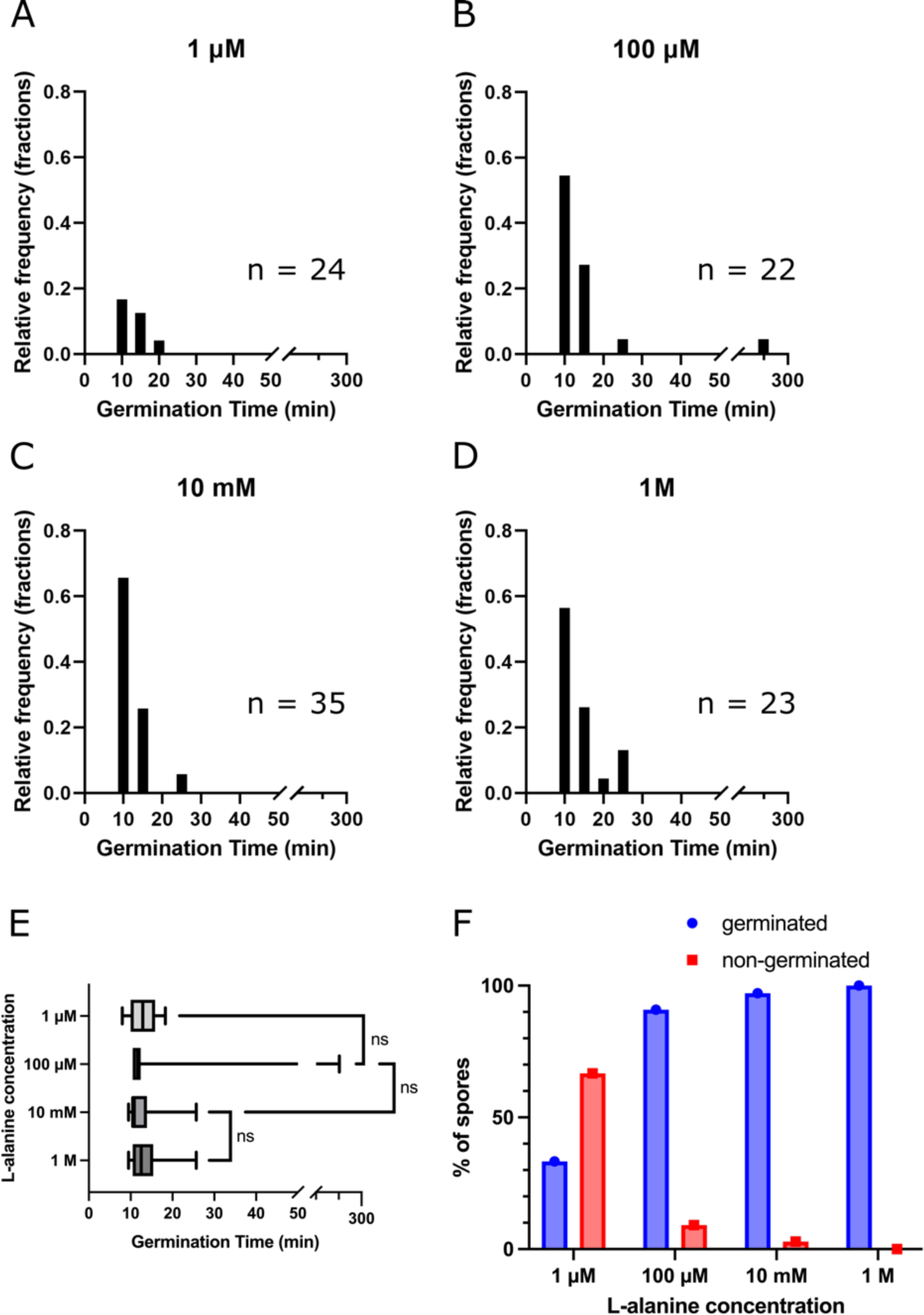
*Bacillus subtilis* germination in response to a concentration gradient of L-alanine. Video 2 contains example timelapses for each condition. **(A-D)** Relative frequency distributions of the time of germination in each condition. The graphs show what proportion (i.e., relative frequency) of the analysed spores germinated within 5-minute intervals. The total number of analysed spores (n) in each condition is indicated. **(E)** Boxplot of the germination times in each condition. The line shows the median, the box extends from the 25^th^ to 75^th^ percentiles and the whiskers from min to max. Mann Whitney tests showed no statistically significant differences (*p* > 0.05) between conditions. **(F)** Bar chart showing the percentage of spores that germinated or remained dormant (non-germinated) after 5 hours of incubation for the four conditions compared within the microfluidic device..

### Responsiveness of *A. oxalaticus* to potassium oxalate

Next, we used the 4CMC device to study germination in the non-model environmental species *A. oxalaticus*, an obligate oxalotrophic endospore-forming species isolated from soil (Estoppey 2023). Oxalic acid and its conjugated base oxalate are abundant in soil and have been shown to be of great importance in ecosystem functioning and interactions amongst species, in particular considering bacterial-fungal interactions (Palmieri *et al*. 2019). Since *A. oxalaticus* is known to be a strict oxalotrophic species (Zaitsev *et al*. 1998), we hypothesised that spore germination should be tightly controlled by the concentration of oxalate in its environment. To better understand the germination of this species, we investigated the responsiveness of *A. oxalaticus* spores upon exposure to a concentration gradient of a germinant, the soluble salt potassium oxalate, ranging from 0.3 g L^-1^ to 8 g L^-1^ (Video 3). Figure 4A shows microscopy images of spore germination. This corresponds to concentrations ranging from 8 g L^-1^ and 4 g L^-1^, to those measured in fungal spent media (0.65 g L^-1^ and 0.3 g L^-1^) (Estoppey *et al*. 2022; Estoppey 2023). At the highest concentration of potassium oxalate (8 g L^-1^), no germination was observed. At 4 g L^-1^, 0.65 g L^-1^ and 0.3 g L^-1^, germination events were observed throughout the 20-hour duration of the experiments, with the last event occurring after 16.6 hours at 0.65 g L^-1^, 19.25 hours at 0.3 g L^-1^ and 16.3 hours at 4 g L^-1^ (Figure 4B-E) Mann Whitney tests showed no statistically significant differences between the different conditions (Figure 4F). The number of spores that germinated after 20 hours was as follows: 0% at 8 g L^-1^, 33.3% at 4 g L^-1^, 56.9% at 0.65 g L^-1^ and 59.4% at 0.3 g L^-1^ (Figure 4G). To investigate the reason behind the lack of germination at high concentrations, the pH of the media was also measured and found to range from 7.0 at 0 g L^-1^ to 7.5 at 8 g L^-1^. We performed an additional control experiment with Schlegel AB + 0.65 g L^-1^ potassium oxalate, Schlegel AB alone (0 g L^-1^ potassium oxalate), NB and physiological water (Video 4 and Supplementary Figure S4)). We observed no germination in NB or physiological water, and only 6.4% germination after 20 hours in Schlegel AB alone, which corresponds to 3 out of 49 analysed spores germinating (Supplementary Figure S4A-E). We found no statistically significant results between the 0.65 g L^-1^ potassium oxalate conditions in the main and control experiments (Mann Whitney test), with median germination times of 295.6 and 297.5 minutes, respectively (Supplementary Figure S4F).

**Figure 4:**
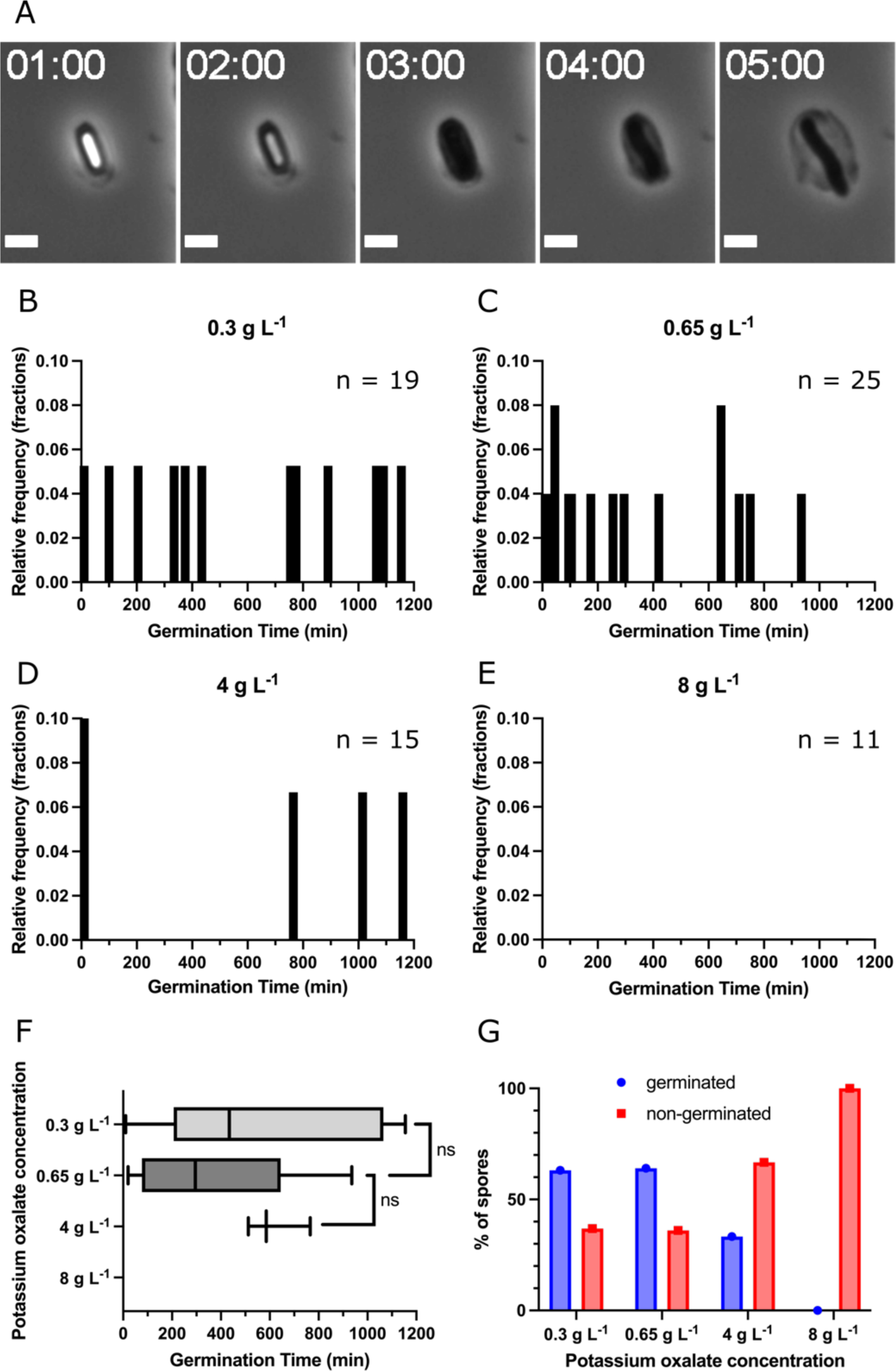
Germination of *Ammoniphilus oxalaticus* in response to a concentration gradient of potassium oxalate. Timelapses for each condition can be found in Video 3. **(A)** Phase contrast microscope images of *A. oxalaticus* showing germination and outgrowth in 0.65 g L^-1^ potassium oxalate. Scale bars, 2 µm. Time stamp format, hh:mm. **(B-E)** Relative frequency distributions of the time of germination in each condition. The graphs show what proportion (i.e., relative frequency) of the analysed spores germinated within 5-minute intervals. The total number of analysed spores (n) in each condition is indicated. **(F)** Boxplot of the germination times in each condition. The line shows the median, the box extends from the 25^th^ to 75^th^ percentiles and the whiskers from min to max. Mann Whitney tests showed no statistically significant differences (*p* > 0.05) between conditions. Note that there is no data for 8 g L^-1^ since there was no germination in that condition. **(G)** Quantification of the spores that remained dormant and germinated after 20 hours of incubation in the microfluidic device in each condition. Results from additional control experiments with Schlegel AB + 0.65 g L^-1^ potassium oxalate, Schlegel AB only (0 g L^-1^ potassium oxalate), NB and physiological water, as well as the comparison between the two Schlegel AB + 0.65 g L^-1^ potassium oxalate conditions (from the main and control experiments) can be found in Supplementary Figure S4, and the corresponding timelapses in Video 4.

### Responsiveness of *T. rossicum* to nutrient conditions

After demonstrating the use of the 4CMC device for both model and non-model soil bacteria, we chose to explore its usability to study soil fungi. There has been a lot of interest in recent years in using *Trichoderma* species as biocontrol agents (Huma 2022; Li *et al*. 2022; Manzar *et al*. 2022; Wang *et al*. 2022; Risoli *et al*. 2023; Sorathiya *et al*. 2023), making it particularly important to understand how *Trichoderma* spores respond to their environment. Here, the responsiveness to nutrient availability was investigated using *T. rossicum*. Spores were exposed to a concentration gradient of rich media (potato dextrose broth, PDB), ranging from 0% (physiological water) to 100% PDB (Video 5). The results are displayed in Figure 5. Figure 5A shows snapshots of live imaging at different time points in each condition. Here, each image spans one complete chamber of the device. We focused on three observations: the swelling of the spores which corresponds to rehydration and surface remodelling (Figure 5B), the time of apparition of the first germ tube (showing germination, Figure 5C) and the diameter of the lead hyphae (Figure 5D). No statistically significant differences (Mann Whitney tests) were observed in the time of apparition of the first germ tube, which was 13.1 ± 2.6 hours at 0.1%, 12.7 ± 3.0 hours at 1% and 12.3 ± 1.3 hours at 100%. At 0% PDB, limited swelling was observed, but no germ tube formation. Both the maximum swelling diameter from the isotropic growth of spores and the diameter of the lead hypha were measured and values were found to be significantly different between each condition. The maximum swelling diameter was 3.5 ± 0.6 µm at 0%, 5.9 ± 1.1 µm at 0.1%, 6.8 ± 1.5 µm at 1% and 8.1 ± 0.9 µm at 100% PDB. The diameter of the lead hypha was 2 ± 0.3 µm at 0.1%, 2.6 ± 0.4 µm at 1% and 3.7 ± 0.8 µm at 100% PDB.

**Figure 5:**
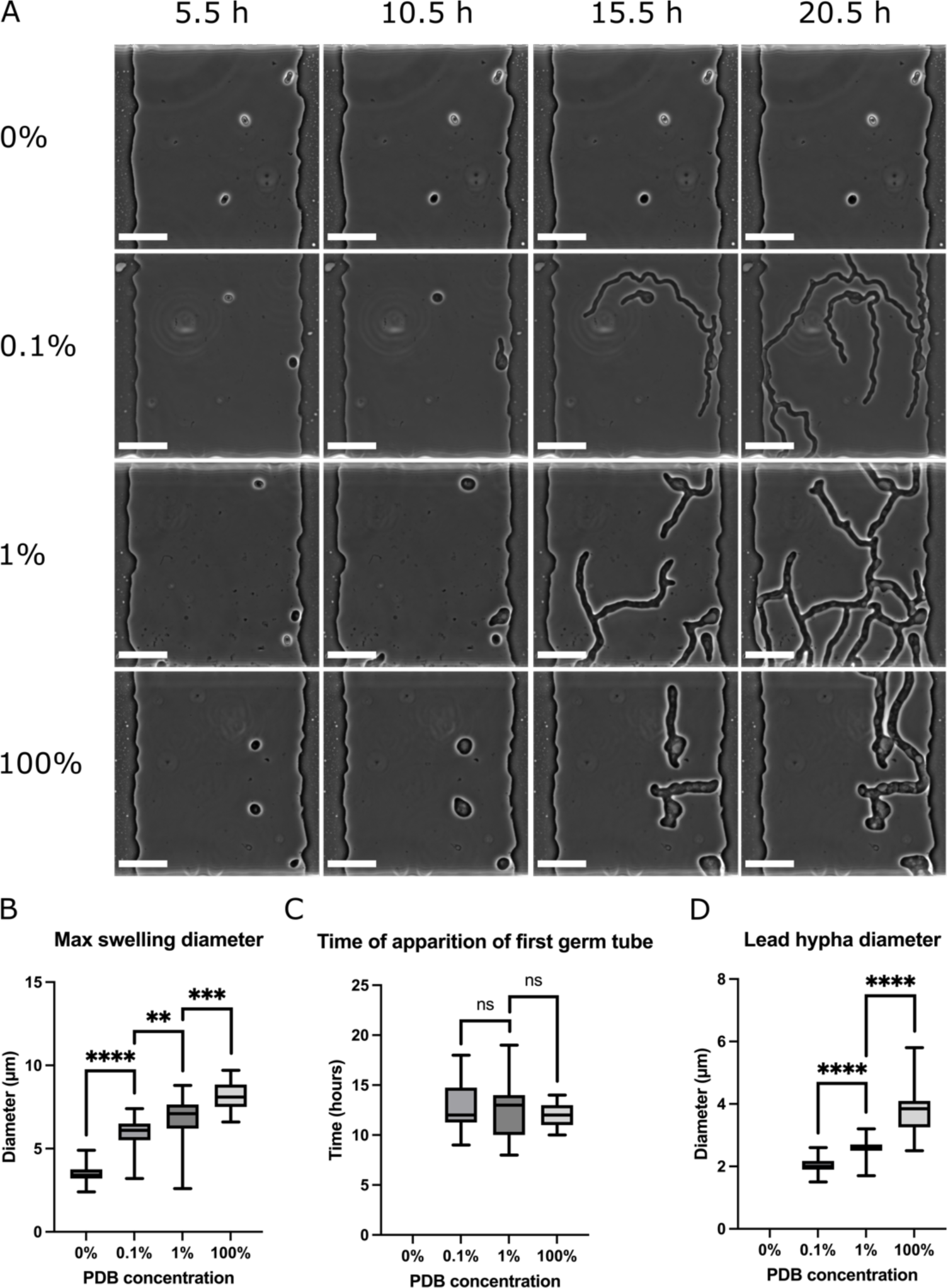
Characterising *Trichoderma rossicum* germination in a concentration gradient of potato dextrose broth (PDB). The number of analysed spores (n) in each condition was the following: 0%: n = 25, 0.1%: n = 19, 1%: n = 22, 100%: n = 25. Timelapses are available in Video 5. **(A)** Snapshots of live imaging of *T. rossicum* germination in each condition. Scale bars, 20 µm. **(B)** Maximum swelling diameter of spores in different PDB concentrations. The line shows the median, the box extends from the 25^th^ to 75^th^ percentiles and the whiskers from min to max. Mann Whitney tests show significant differences (p < 0.05) between each condition. **(C)** Time of apparition of the first germ tube at different PDB concentrations. The line shows the median, the box extends from the 25^th^ to 75^th^ percentiles and the whiskers from min to max. No significant differences (p > 0.05) were found according to Mann Whitney tests. **(D)** Diameter of the lead hypha in different PDB concentrations. The line shows the median, the box extends from the 25^th^ to 75^th^ percentiles and the whiskers from min to max. Mann Whitney tests show significant differences (p < 0.05) between each condition.

## Discussion

The aim of this study was to design and utilise a microfluidic platform for the investigation of spore germination in multiple conditions simultaneously, applicable for model and non-model bacterial and fungal spores. Our design builds on previous work by Barkal *et al*. (Barkal *et al*. 2016), but bringing all four conditions together into one device. This allows for streamlining of the loading process and imaging of multiple conditions at high magnification (where the use of oil limits the area that can be covered). The former is important since the time between loading and the start of the experiment should be kept to a minimum to avoid cells adhering to the device outside chambers (to avoid clogging the supply channels), and to allow to start recording as fast as possible (as germination can start quickly upon exposure to the germinant). The device was developed to be easy to use, reliable and effective. Besides the microscope, running experiments requires little specialist equipment: tubing and connectors, syringes, and a syringe pump. It is compatible with existing image analysis software (such as ImageJ with the SporeTrackerX plugin (Omardien *et al*. 2018)), and highly versatile.

We initially characterised the device by using it to quantify the germination of *B. subtilis* in rich media (NB), and synthetic media (SM) with and without the addition of L- or D-alanine (at 10 mM concentrations). Alanine was used in this initial experiment as it acts as a germinant in its L-enantiomeric form but has an inhibitory effect in its D-form (Yasuda and Tochikubo 1985). As expected, we observed little germination with D-alanine (7.4%) or SM alone (4.4%), and full germination (100%) in L-alanine and a rich medium such as NB. It is important to note here that bacterial spore germination studies vary greatly in measurement methods and strains used, but also in how spores are prepared and stored, as well as how germination is induced. This makes it difficult to compare results across different studies. Nevertheless, the results obtained here are in agreement with previous studies. For instance, our results are consistent with the flow cytometry measurements of 90% of germinant cells after 20 minutes obtained in a study investigating the effect of the spore’s coat protein CotG in different *B. subtilis* strains in conditions close to the SM + L-alanine condition used in the present study (Saggese *et al*. 2022). Indeed, we measured 79% germination after 20 minutes, using a different strain and without heat activation, a process used to speed up and increase germination (Luu *et al*. 2015). Cangiano *et al*. had used similar experimental conditions and measured just over 80% germination after 20 minutes with flow cytometry with the same wild type strain (PY79) (Cangiano *et al*. 2014). As with flow cytometry, our method provides single-cell level information, but it has the additional advantage of providing continuous data over time. By looking at the germination times of each individual spore, we observed more heterogeneity (a wider distribution of germination times) in SM + L-alanine compared to rich media (NB). Wu *et al*. used phase-contrast microscopy together with Raman tweezers to specifically investigate the heterogeneity of cell lysis of *B. subtilis* cells during spore germination, upon prophage induction (Wu *et al*. 2021). They found cell elongation to start after around 100 minutes in rich media. While we did not measure cell elongation itself, this is in line with our observations.

The next set of experiments consisted of investigating germination at various concentrations of L-alanine in more detail. The pathways involved in endospore germination have been extensively studied in the past few years and summarised in detail by Lyu *et al*. (Lyu *et al*. 2023). Briefly, receptors located in the inner membrane or cortex receive signals from germinants, eventually triggering germination. Enough of the appropriate germinants must be present for their interactions with germinant receptors to reach a certain threshold and therefore lead to a cascade of events, including the release of DPA and rehydration of the spore’s core. In the case of the germinant L-alanine, it binds to and activates the germinant receptor GerA, leading to germination. The heterogeneity of germination times is thought to be due to differences in levels of GerA in each spore. Similarly, there could be differences in levels of the other components of the pathway downstream of this initial event. In our experiment there were no statistically significant differences in the times of germination for the concentrations measured. Still, as expected, the overall germination increased with increasing concentration of germinant, from 33.3% at 1 µM to 90.9% at 100 µM, 97.1% at 10 mM, and 100% at 1 M concentration. Lu *et al*. studied in detail the effect of L-alanine on different wild type and lab strains of by measuring DPA release, showing significant differences among different strains (Lu *et al*. 2021). Upon further analysis of the strains with higher germination potential in response to L-alanine, they found that at least 100 µM concentration was required to induce spore germination (in wild type strains), whereas we did observe some germination at 1 µM already. In another strain (PY79), Krajčíková *et al*. found little germination (around 20% loss of OD after 60 minutes) at 10 µM and much higher germination (over 50% loss of OD after 60 minutes) at 1 mM L-alanine, which is in line with our observations (Krajčíková, Bugárová and Barák 2021). For the PS533 strain, Butzin *et al*. analysed germination by measuring DPA release for L-alanine concentrations ranging from 500 µM to 1 M (Butzin *et al*. 2012). Contrary to our measurements, they observed a decrease in germination as the concentration increased, with 75% of germination at 1 M after 2 hours, just over 75% germination after 2 hours at 500 µM, and around 90% germination after 2 hours for all the concentrations in between. They hypothesise that the decrease of germination at high concentration could be due to nonspecific inhibition caused by high ionic strength, for example. Frentz and Dworkin studied *B. subtilis* germination heterogeneity in various concentrations of L-alanine using a bacterial luminescence system to measure differences in metabolic activity (Frentz and Dworkin 2020). Unlike us, they observed no spore germination at a concentration of 1 µM with their experimental conditions. These results further demonstrate the difficulty to compare spore germination studies due to the high variations in methods.

In addition to germinants, other molecules (e.g., DPA) can also induce germination by activating intermediate steps. This could explain the correlation between spore density and germination times that was observed in rich media, as molecules released by the first spores that germinate could contribute to the germination of neighbouring spores. Evidence that spore density influences germination has been found before. For example, Caipo *et al*. showed that higher concentrations of *Bacillus megaterium* spores resulted in faster germination as well as higher overall percentage of germination, which could be a result of quorum sensing or chemical signalling between spores (Caipo *et al*. 2002). Webb *et al*. on the other hand, only found local spore density to have a minor influence on germination of spores of *Clostridium botulinum* compared to the effect of the total number of spores in the experiment (Webb *et al*. 2012). However, those experiments were performed on agar slides, where the local growth conditions are difficult to control and define. Microfluidic chemostat systems provide an alternative format, where small chambers with controlled conditions can be analysed. In our microfluidic device, we found a negative correlation (r = -0.8) between the number of spores per chamber and their germination time in rich media. Interestingly, we did not see a correlation in the SM + L-alanine condition, where spores are able to germinate but not to grow (spores turn phase-dark but there is no outgrowth and elongation). This suggests that in this case, the cascade of events triggered by exposure to the germinant does not progress far enough to release the molecules needed to accelerate the germination of neighbouring spores.

*Bacillus* spores are the most studied example of endospore germination, followed by *Clostridium* spores. In both cases, most observations stem from model lab strains. However, very little is known about other species or strains, and there might be different mechanisms at play for environmental species. We studied the obligate oxalotrophic endospore-forming bacterium *A. oxalaticus,* which was isolated from a soil sample (Estoppey 2023). Very little is known about this species, previously described by Zaitsev *et al*. (Zaitsev *et al*. 1998). Oxalic acid is a ubiquitous compound produced by fungi, bacteria, plants and animals and plays important roles in varied ecosystem functions (Palmieri *et al*. 2019). We compared different concentrations of potassium oxalate, a water-soluble compound formed by potassium and the conjugated base of oxalic acid oxalate. The concentrations tested ranged from concentrations corresponding to those measured in fungal exudates (0.3 and 0. 65 g L^-1^) to those used under laboratory conditions to grow oxalotrophic microorganisms (4 and 8 g L^-1^) (Schilling and Jellison 2004; Estoppey *et al*. 2022; Estoppey 2023). As expected, we observed germination at 0.3 and 0.65 g L^-1^ (59.4% and 56.9% after 20 hours, respectively). However, we found little germination at 4 g L^-1^ and none at 8 g L^-1^. We measured the pH of the media used for the experiment (Schlegel AB) in order to assess whether this was a pH-mediated effect, given that oxalic acid is a strong organic acid (pKas 1.25 and 4.28). However, since the medium was buffered, the pH only increased from 7 to 7.5 at the highest potassium oxalate concentration. Hence, we hypothesise that the lack of growth at the highest potassium oxalate concentration could be due to the concomitant presence of oxalic acid/oxalate and potassium, both potentially inducing osmotic stress.

The final set of experiments consisted of observing germination of the fungus *T. rossicum* in a concentration gradient of PDB media. While there are numerous germination studies in bacteria (and particularly in endospore-forming bacteria), much less is known about fungal spore germination. However, fungal spores can be used in many applications, such as in the formulation of biocontrol agents, many of which contain *Trichoderma* spores (Martinez *et al*. 2023). Biocontrol formulations should maintain their functional properties over time to allow for long shelf lives. Hence, spore germination should be understood so it can be prevented during storage. Spore germination understanding is also required to ensure germination and effective biocontrol once the formulation is applied to soil, overcoming soil fungistasis while limiting undesired side-effects (Garbeva *et al*. 2011; Massart, Margarita and Jijakli 2015). Our data indicate that as nutrient availability decreases, the time of germination remains the same; the swelling of the spores and the diameter of hyphae, however, are reduced, which has not been shown before to the best of our knowledge. This could be a way to optimise the ratio between nutrients available for growth and the expansion of the mycelium. Indeed, the concept of adaptive foraging strategies has been explored in fungi, demonstrating the ability to exploit new resources and redirect efforts to that location (Tlalka *et al*. 2008). Tradeoffs in mycelial architecture, such as between branching (represented by internodal length) and hyphal diameter have also been described (Lehmann *et al*. 2019). Microfluidic technologies have been used previously to investigate fungal foraging behaviour and hyphal space exploration (Aleklett *et al*. 2021). Additionally, there is evidence that fungal biomass allocation depends on their growth rate (Veresoglou *et al*. 2018). However, very little is known about those processes at the time of germination as studies usually focus on established mycelium. Our data suggest that resource allocation is already determinant at the time of spore germination.

## Conclusion

We presented here the 4CMC, a microfluidic platform for spore germination experiments with model and non-model bacterial and small fungal spores. We focused on spore-forming soil dwellers to better our understanding of their germination, which is required to be able to use spores for biotechnological applications such as biocontrol agent formulations. Our observations indicate that higher concentrations of *B. subtilis* spores result in faster germination in rich media. We studied the response of the environmental species *A. oxalaticus* to various concentrations of potassium oxalate, providing the first quantitative, high temporal and spatial resolution information regarding the germination of this obligate oxalotroph, and showing that low concentrations of potassium oxalate (that correspond to concentrations measured in fungal exudates) result in more spores germinating as compared to higher concentrations used under laboratory conditions. The 4CMC will allow to investigate other sources of oxalate for oxalotrophic bacteria and could help understand their interactions with other soil dwellers, in particular with oxalogenic fungi. Additionally, our results demonstrate for the first time that the diameter of *T. rossicum* spores and hyphae increases with increased nutrient concentration, while the germination time remains unchanged. Our fungal spore study only explored spore swelling, germ tube apparition and hyphal diameter, but further studies using this device could investigate the effect of spore density, hyphal branching, or even chlamydospore formation and germination. This device would also be well suited to study communities or consortia, in various trophic and abiotic conditions that can be manipulated in a solution. As a result, the 4CMC allows exploring at a microscale a combination of biotic, trophic, and abiotic factors, making it a relevant tool to understand microbial behaviour from an environmental perspective but under controlled conditions. It can also be adapted to different types of responsiveness experiments beyond spore germination. For example, to evaluate the effect of inducers on engineered cells. Finally, the flexibility of the PDMS allows for a small range of sizes of spores (e.g. both conidia and chlamydospores of *T. rossicum*) to be studied with the current design, but it can be adapted to accommodate larger organisms/spores by changing the height of the microchambers.

## Author contributions

The idea for the 4CMC platform and study was conceived by C.E.S and L.S.B. L.S.B designed and fabricated the microfluidic device, and performed the experiments, the microscopy imaging and the data analysis. A.E. isolated and provided *A. oxalaticus* and performed initial proof-of-concept experiments. C.E.S supervised this study, whereas S.B, G.-B.S. and P.J. provided valuable feedback to the experimental work. L.S.B. and C.E.S took the lead in writing the manuscript with contributions from all other authors. All authors read and approved the final version of the manuscript.

## Acknowledgements

We acknowledge financial support from the Department of Bioengineering at Imperial College London, as well as Kate Collins for generating the COMSOL simulation.

