## Supplementary Information for "Microfluidic platform for microbial spore germination studies in multiple growth conditions"

### Supplementary materials

A

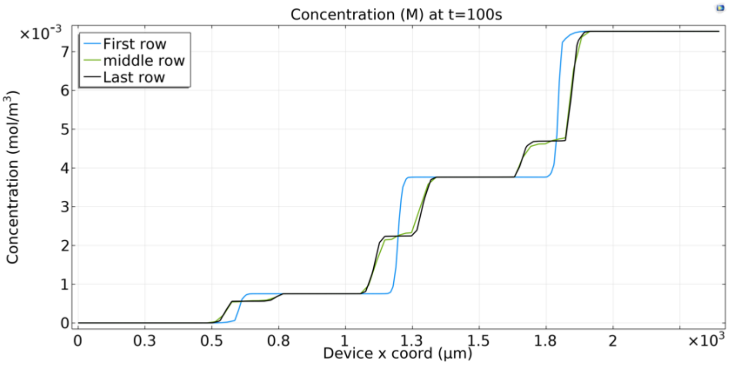


B

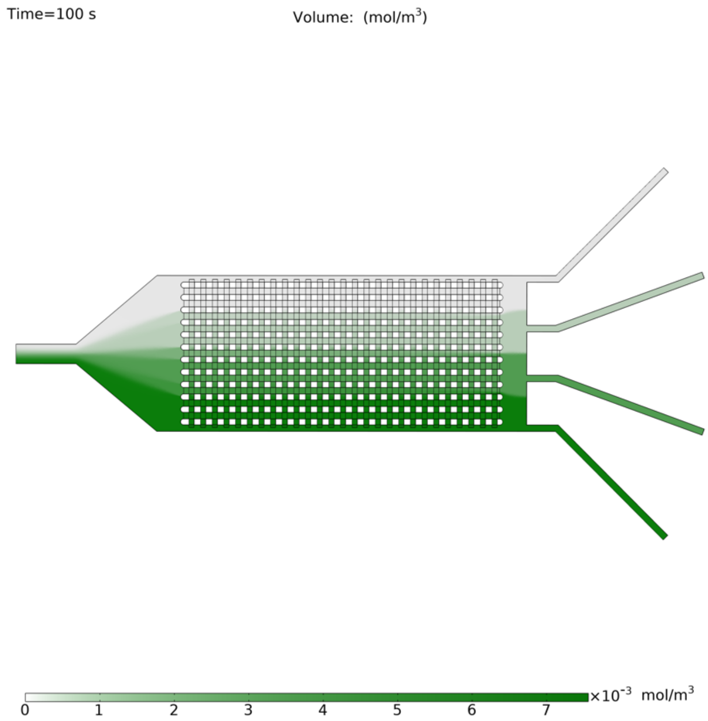


**Figure S1: COMSOL simulations.** Throughout our COMSOL simulations, the fluorescein diffusion coefficient was set to $0.4 \times 10^{-5}$ $cm^2$ $s^{-1}$ (Casalini et al., 2011). **(A)** Concentration of fluorescein across the device (spanning each zone corresponding to each concentration) in the first row (on the left side on B), one in the middle and the last one (on the right side on B), showing our 4 distinct concentrations are maintained across the length of the device. **(B)** Fluorescein concentrations in the device.

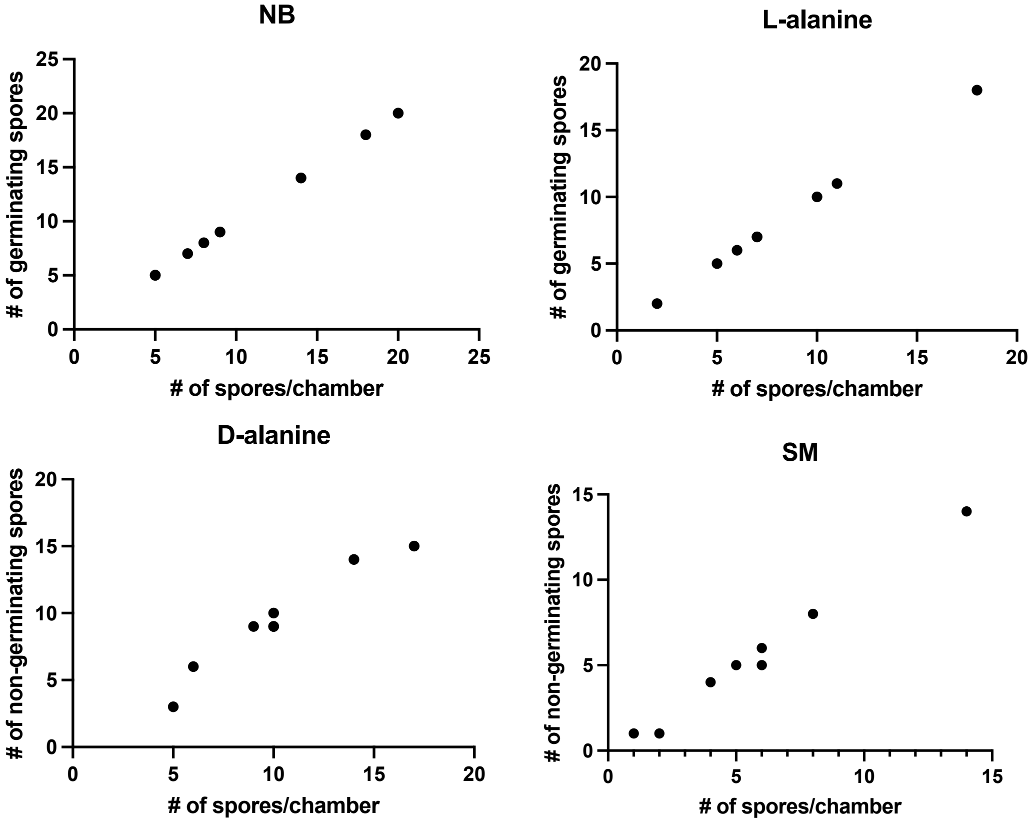


**Figure S2: Number of spores germinating (or not germinating) in relation to number of spores per chamber.** In those four conditions, the number of spores in a chamber is strongly correlated (Spearman's rank correlation) to the number of spores germinating (NB r = 1, L-alanine r = 1), or non-germinating (D-alanine r = 0.95, SM r = 0.98).

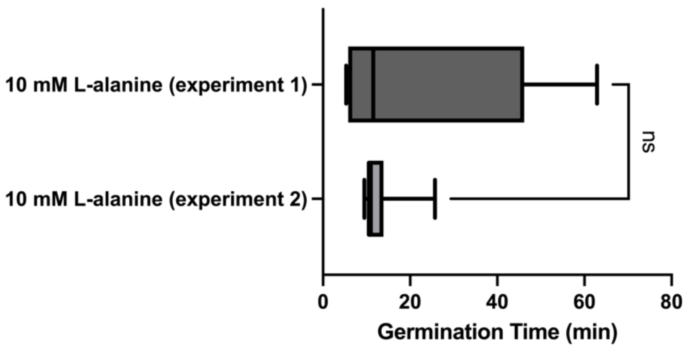


**Figure S3: Comparison of germination times in the two 10 mM L-alanine conditions.** Boxplot of the germination times in the 10 mM L-alanine conditions from both sets of experiments. The line shows the median, the box extends from the 25th to 75th percentiles and the whiskers from min to max. A Mann Whitney test showed no statistically significant differences ($p$ > 0.05) between the two conditions.

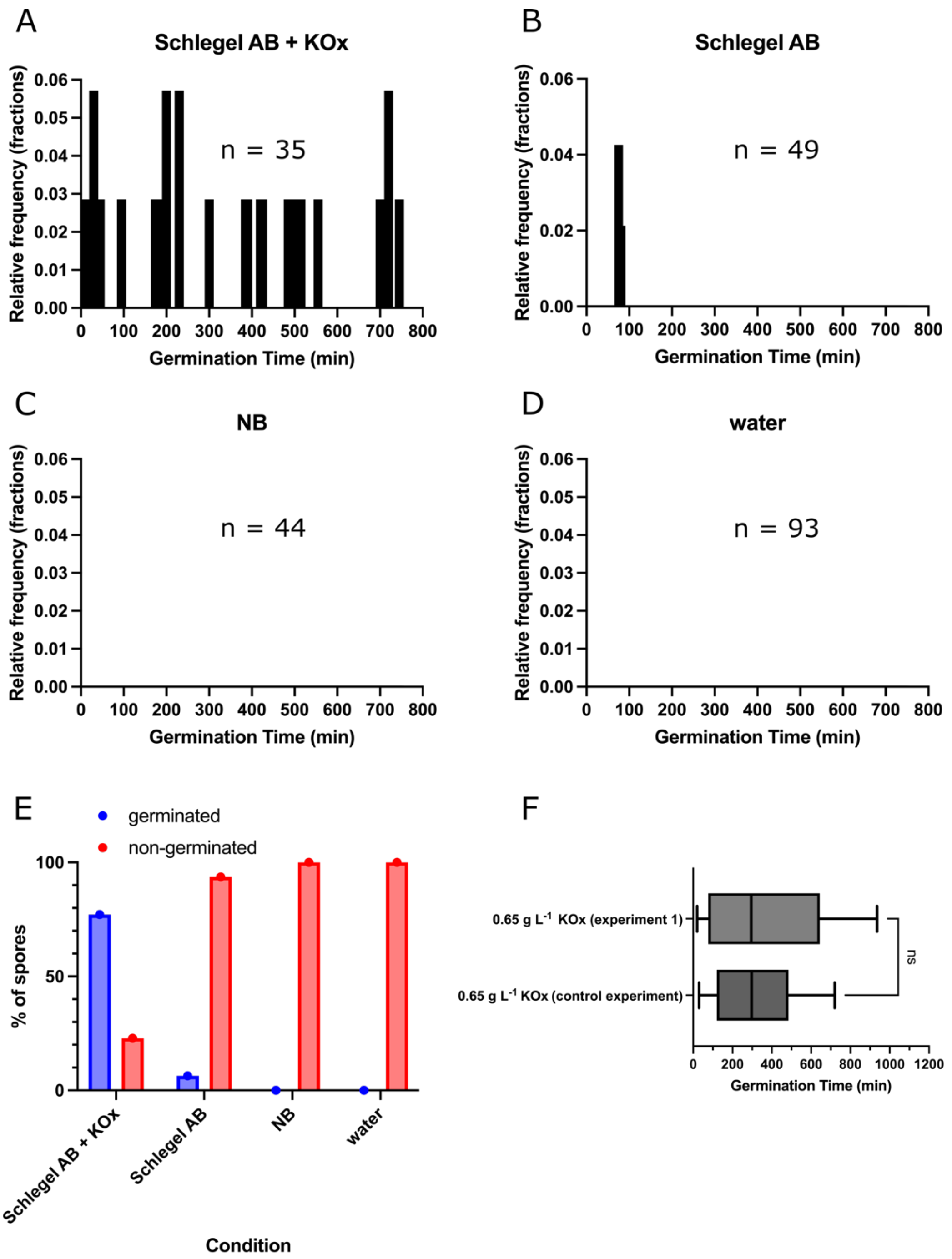


**Figure S4: Germination of *Ammoniphilus oxalaticus* in 4 conditions. (A-D)** Relative frequency distributions of the time of germination in each condition: Schlegel AB + 0.65 g $L^{-1}$ potassium oxalate, Schlegel AB (0 g $L^{-1}$ potassium oxalate), NB and physiological water. These graphs show what proportion (i.e., relative frequency) of the analysed spores germinated within each 5-minute interval. The number of analysed spores (n) in each condition is indicated. **(E)** Bar chart showing the percentage of spores that germinated or remained dormant (non-germinated) after exposure to each condition for 20 hours within the microfluidic device. **(F)** Boxplot of the germination times in the Schlegel AB + 0.65 g $L^{-1}$ potassium oxalate conditions from the main and control experiments. The line shows the median, the box extends from the 25$^{th}$ to 75$^{th}$ percentiles and the whiskers from min to max. A Mann Whitney test showed no statistically significant differences ($p > 0.05$) between the two conditions.

**Videos**

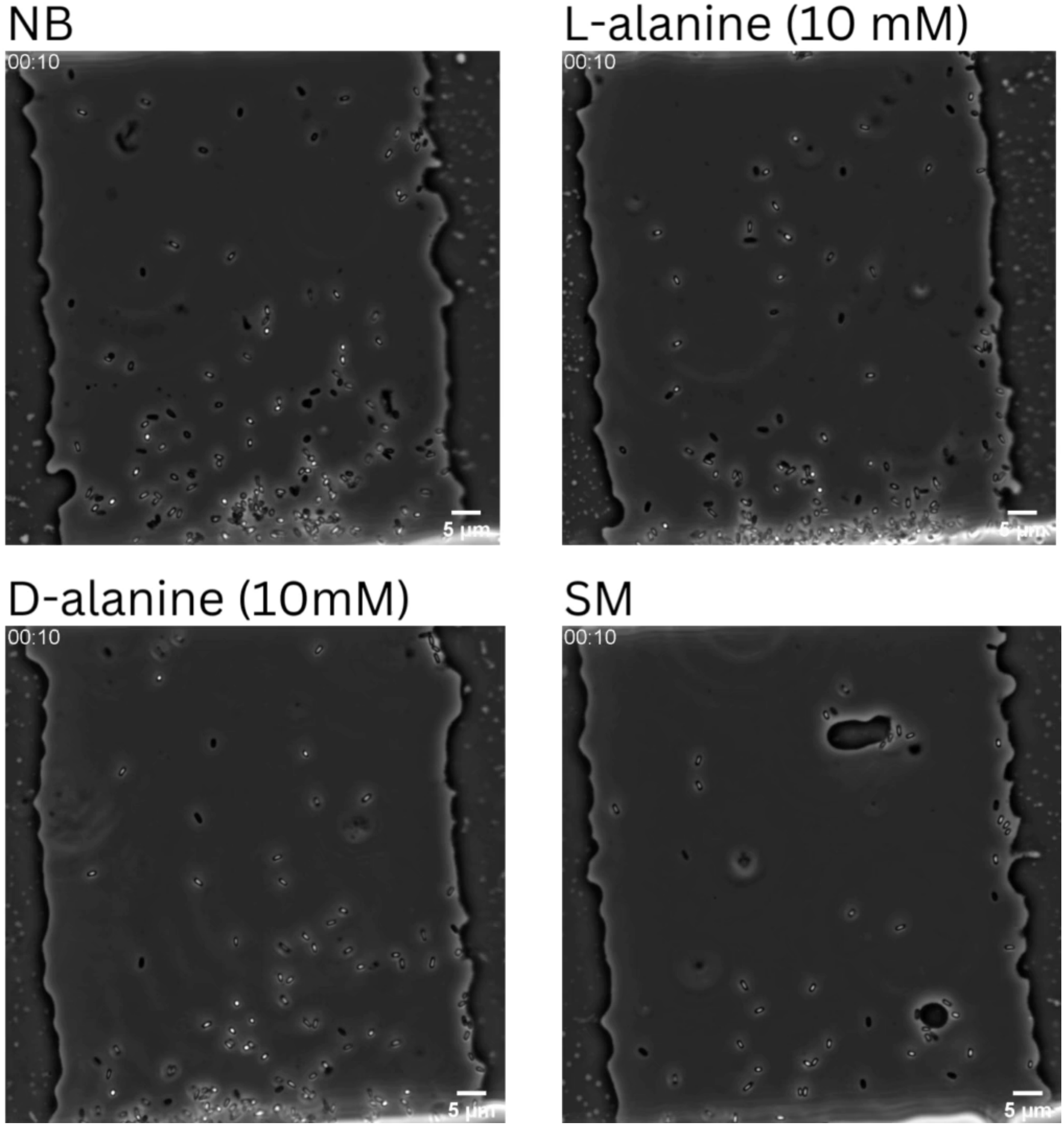


**Video 1: Timelapses of germination of *Bacillus subtilis* in the microfluidic device over 5 hours in 4 conditions.** Time stamp format is hh:mm.

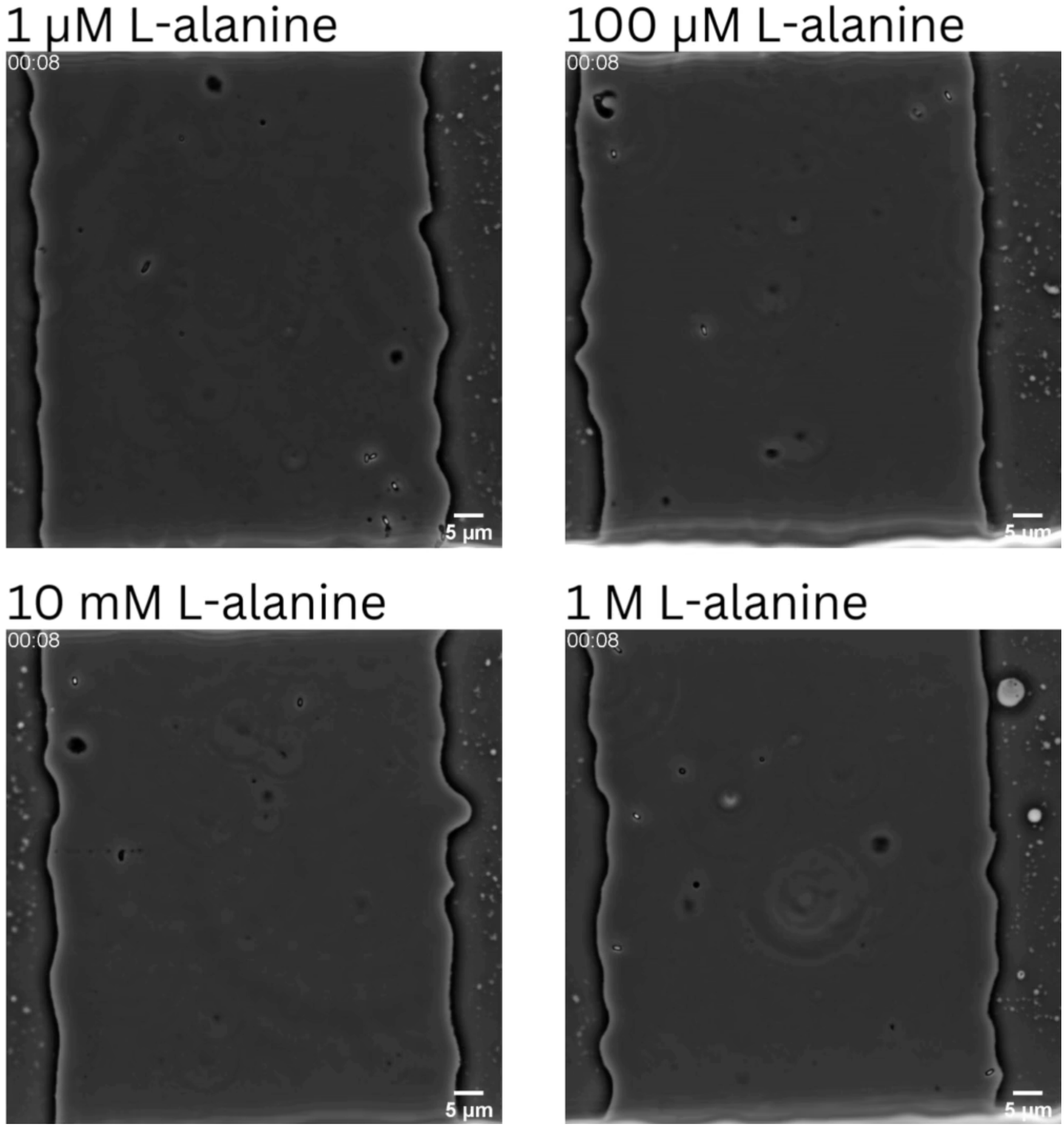


**Video 2: Timelapses of germination of *Bacillus subtilis* in the microfluidic device over 5 hours in a concentration gradient of the germinant L-alanine.** Time stamp format is hh:mm.

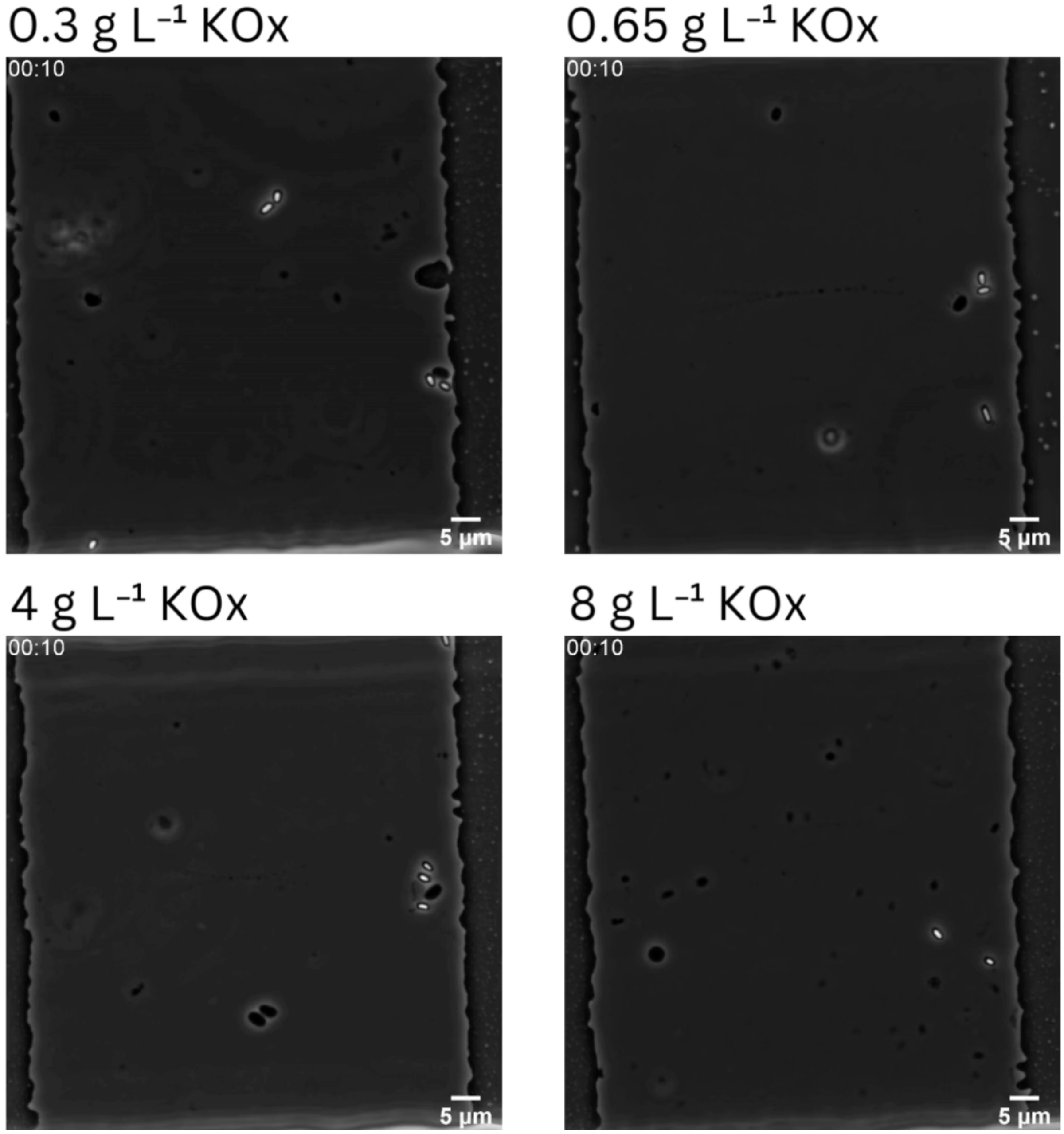


**Video 3: Timelapses of germination of *Ammoniphilus oxalaticus* in the microfluidic device over 20 hours in a concentration gradient of the germinant potassium oxalate.** Time stamp format is hh:mm.

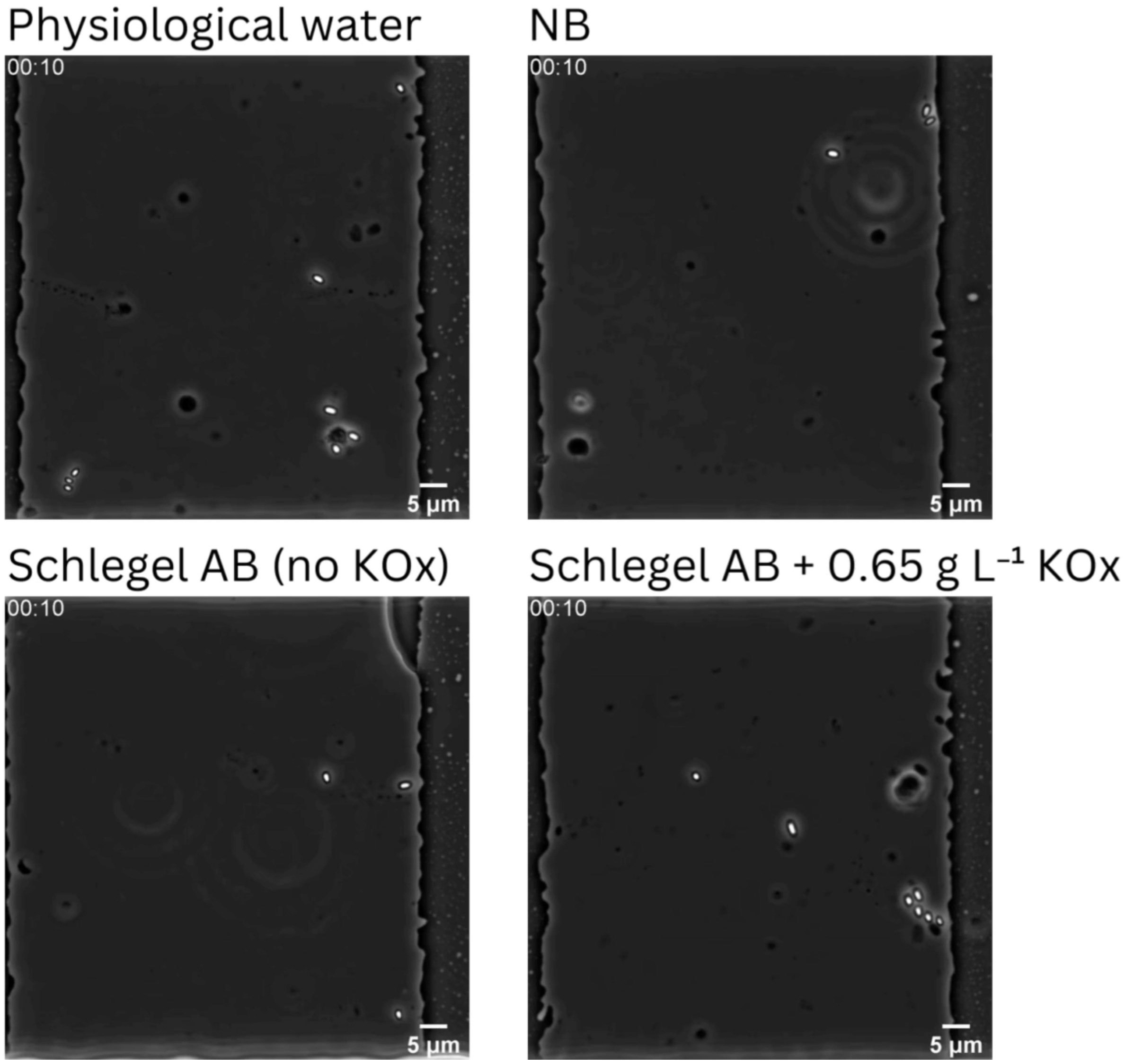


**Video 4: Timelapses of germination of *Ammoniphilus oxalaticus* in the microfluidic device over 20 hours in 4 different conditions.** Time stamp format is hh:mm.

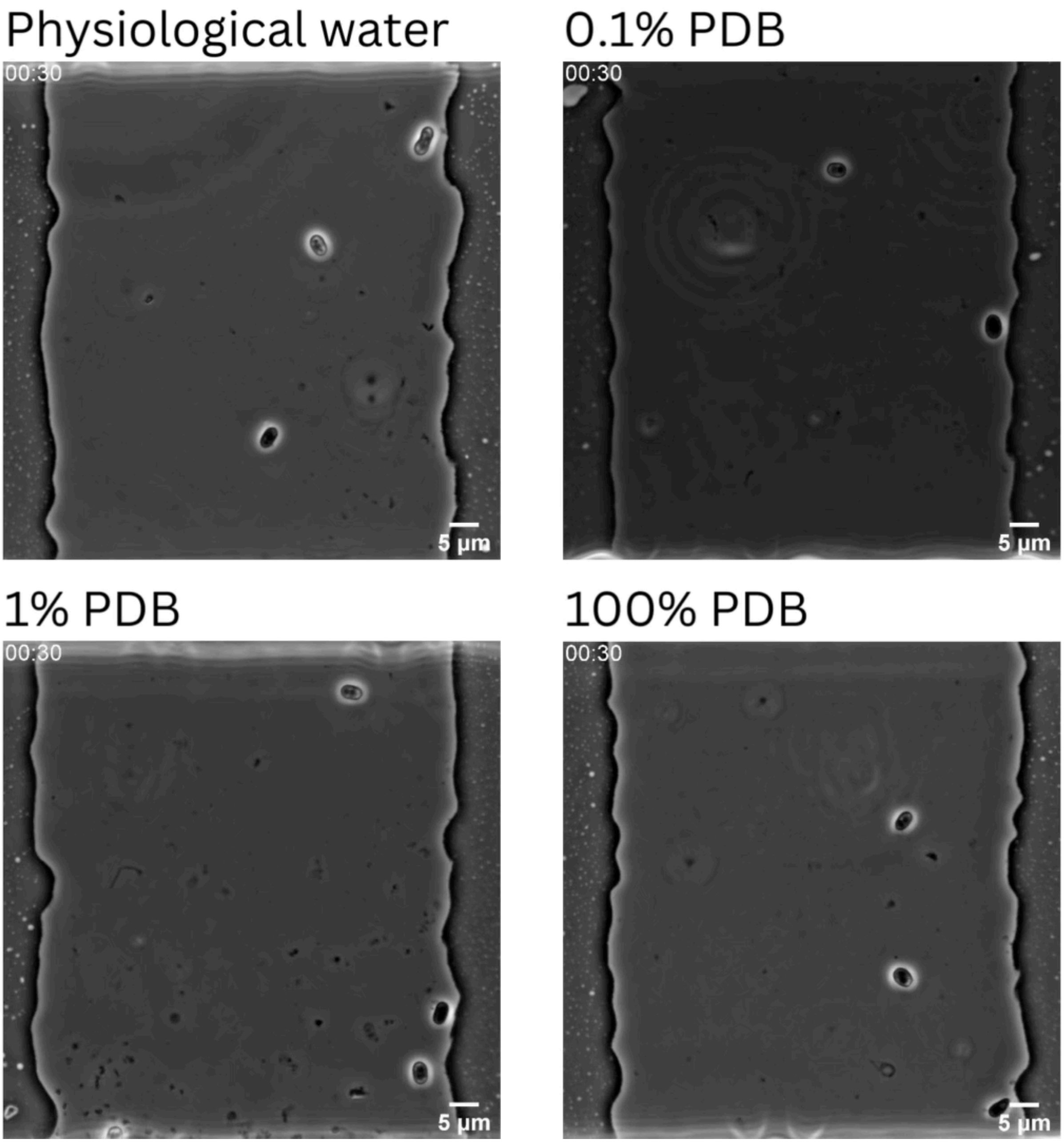


**Video 5: Timelapses of germination of *Trichodrema rossicum* in the microfluidic device over 20 hours in a concentration gradient of PDB.** Time stamp format is hh:mm.
